# Light-Sheet Fluorescence Imaging Charts the Gastrula Origin of Vascular Endothelial Cells in Early Zebrafish Embryos

**DOI:** 10.1101/2020.05.27.118661

**Authors:** Meijun Pang, Linlu Bai, Weijian Zong, Xu Wang, Ye Bu, Connie Xiong, Jiyuan Zheng, Jieyi Li, Weizheng Gao, Zhiheng Feng, Liangyi Chen, Jue Zhang, Heping Cheng, Xiaojun Zhu, Jing-Wei Xiong

## Abstract

It remains challenging to construct a complete cell lineage map of the origin of vascular endothelial cells in any vertebrate embryo. Here, we report the application of *in toto* light-sheet fluorescence imaging of embryos to tracing the origin of vascular endothelial cells (ECs) at single-cell resolution in zebrafish. We first adapted a previously-reported method to mount embryos and light-sheet imaging, created an <u>a</u>lignment, <u>f</u>usion, and <u>e</u>xtraction all-<u>i</u>n-<u>o</u>ne software (AFEIO) for processing big data, and performed quantitative analysis of cell lineage relationships using commercially-available Imaris software. Our data revealed that vascular ECs originated from broad regions of the gastrula along the dorsal-ventral and anterior-posterior axes, of which the dorsal-anterior cells contributed to cerebral ECs, the dorsal-lateral cells to anterior trunk ECs, and the ventral-lateral cells to posterior trunk and tail ECs. Therefore, this work, to our knowledge, charts the first comprehensive map of the gastrula origin of vascular ECs in zebrafish, and has potential applications for studying the origin of any embryonic organs in zebrafish and other model organisms.

## Introduction

Determining cell lineage relationships is one of the most fundamental questions in the fields of developmental biology and genetics. An elegant study has charted the lineage tree of *Caenorhabditis elegans* with all the cleavages from the one-cell stage to the adult worm (Sulston et al., 1983). Cell lineage tracing methods fall into two major categories: optical tracing by dye injection, transgenic reporters, or tissue-specific genetic recombination reporters, and sequencebased tracing by either viral barcodes or genome-editing barcodes with CRISPR/gRNA (Spanjaard and Junker, 2017; Woodworth et al., 2017). Due to the regulatory complexity of the lineages and intermingling cell divisions and migration, it remains challenging to apply these methods to the construction of complete cell lineage trees of the organs of early embryos at single-cell resolution in any vertebrate species.

In zebrafish embryos, as in other vertebrates, hematopoietic and endothelial cells (ECs) arise in close association and are thought to be derived from the ventral mesoderm (Detrich et al., 1995; Gore et al., 2012). Single-cell resolution fate maps of the late blastula and gastrula with 2,3-dimethyl 2,3-dinitrobutane-caged fluorescein dextran provided the very first *in vivo* evidence that individual cells give rise to both hematopoietic cells and ECs, and that the majority of ECs are derived from the ventral-lateral region of the shield embryo in zebrafish (Vogeli et al., 2006). However, a comprehensive lineage map of the origin of vascular ECs in vertebrates has not yet been reported.

Advanced optical live-imaging methods have shed new light on approaching this goal; among these, light-sheet fluorescence microscopy (LSFM) has advantages such as low photo-toxicity, rapid imaging, and a capacity for long-term three-dimensional imaging (Huisken and Stainier, 2009). *In toto* imaging of the early stages of fly, zebrafish, and mouse embryos at single-cell resolution has now been reported (Huisken and Stainier, 2009; Keller et al., 2008; McDole et al., 2018). Recently, LSFM has been successfully applied to documenting neuronal cell lineages, movements, and activities in the entire spinal cord of live zebrafish embryos (Wan et al., 2019). Although it often has superior performance, LSFM is not as well used as traditional imaging methods because of the “do-it-yourself ethic” and the problem of big data (Power and Huisken, 2017). In this study, we used LSFM to acquire and resolve *in toto* images of early zebrafish embryos, and charted a comprehensive origin map of all vascular ECs at single-cell resolution – this was made possible by a combination of Zeiss Z.1 LSFM (Carl Zeiss, Jena, Germany), commercially-available Imaris software, and the user-friendly image alignment, fusion, and extraction all-in-one (AFEIO) software that we created. Furthermore, this LSFM system together with the AFEIO software makes it possible to construct the cell-lineage trees of other organs in zebrafish and other model organisms.

## Results

### Optimized mounting and dual-view light-sheet microscopy imaging

To establish the gastrula origin of vascular ECs, we took advantage of LSFM to retrieve full coverage of all developmental cell divisions and migrations in Tg(*H2A.F/Z*:EGFP) transgenic embryos (Huisken and Stainier, 2009; Keller et al., 2008) along with the Tg(*kdrl*:mCherry) transgenic reporter for locating vascular endothelial progenitors/cells. Tg(*H2A.F/Z*:EGFP)^tg/+^; Tg(*kdrl*:mCherry)^tg/+^ double transgenic embryos were used for all imaging experiments (Fig. 1). It has been reported that specimen rotation combined with multiview imaging decreases the degradation of imaging quality induced by tissue scattering and absorption (Swoger et al., 2007), as well as improving the axial optical resolution by compositing with an isotropic point-spread-function (Krzic et al., 2012). On the other hand, specimen rotation requires high mechanical stability, which can be achieved by using a 1% or higher agarose matrix. A previous study reported that 0.4% or higher agarose interferes with embryo growth and mobility while fluorinated ethylene propylene (FEP) with low concentrations of agarose is both optically clear and sufficiently confines the living embryo in a physiological environment, thus providing the optimal choice for live imaging (Kaufmann et al., 2012). Early zebrafish embryos frequently twitch in the absence of tricaine when they are mounted with their intact chorion inside an E3-filled FEP tube (Weber et al., 2014). However, prolonged exposure to tricaine suppresses the contraction of cardiac, skeletal, and smooth muscles and thus affects the hemodynamics (Culver and Dickinson, 2010; Muntean et al., 2010). To decrease the effect of both the chorion and tricaine, we optimized the mounting method by embedding dechorionated embryos in 0.2% low-melting-point agarose in FEP to guarantee both mechanical stability and normal embryonic development, as well as to ensure the exchange of oxygen and fluids during imaging. Tricaine and other drugs, if needed, could be added to the E3 medium at any time (Fig. 1).

**Fig. 1.**
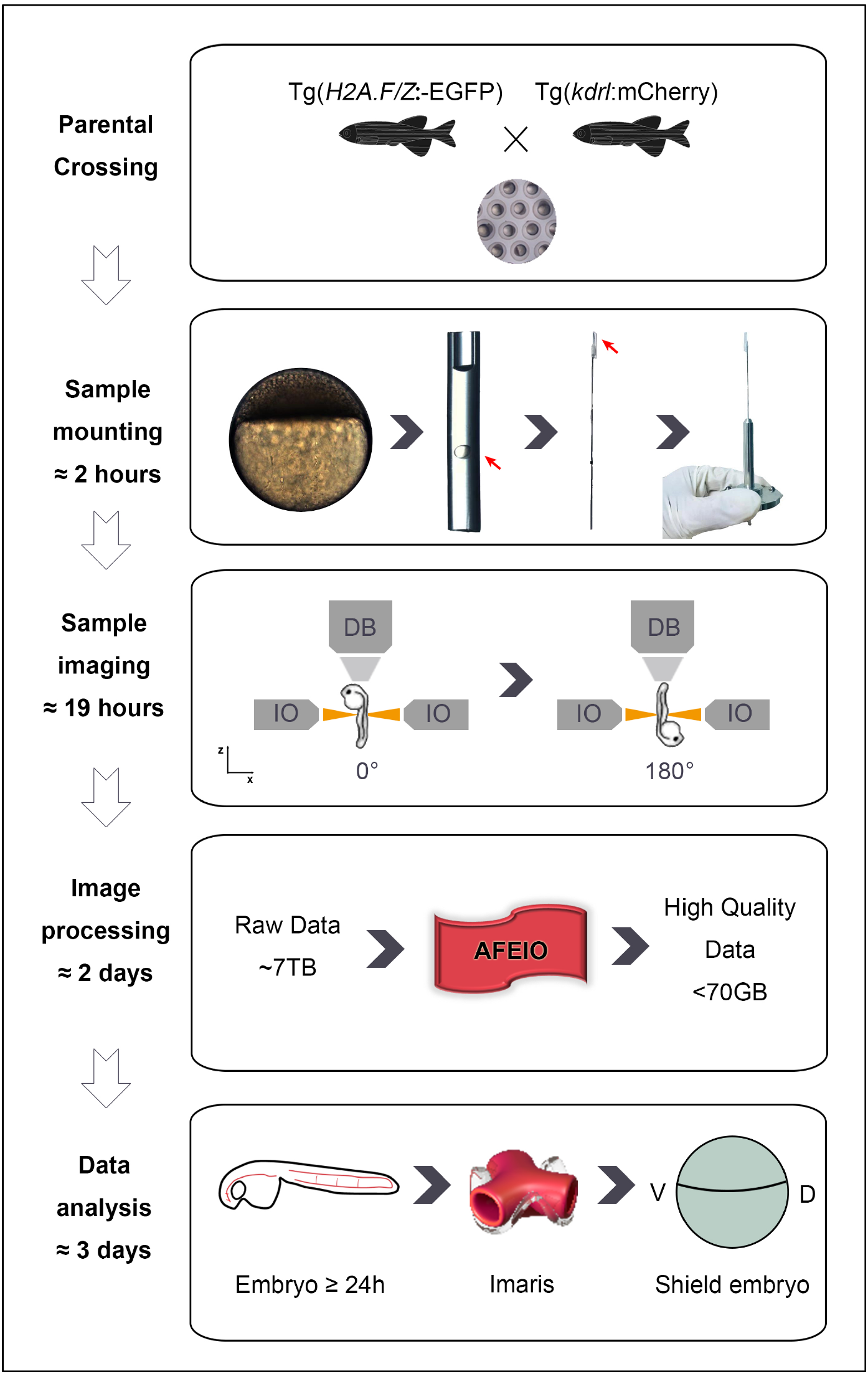
Flowchart of high-resolution imaging of zebrafish embryos with a Zeiss light-sheet fluorescence microscope. **Parental Crossing:** collection of transgenic embryos from crosses between Tg(*H2A.F/Z*:EGFP) and Tg(*kdrl*:mCherry) transgenic lines. **Sample mounting:** careful de-chorionation of embryos at 4 hpf and their transfer into a fluorinated ethylene propylene tube filled with 0.2% agarose. The tube is fixed to a fine wire and then mounted to a holder. **Sample imaging:** The illumination objectives (IO) illuminate the sample from the left and right alternately, while the detection objective (DB) detects signals at 0°. The embryo is then rotated and the 180° images are acquired in the same way. **Image processing:** The raw data (~7 TB) are processed using the ‘alignment fusion and extraction all-in-one (AFEIO)’ software to obtain fused high-resolution data (~70 GB). **Data analysis:** The processed data are imported into Imaris software to run retrospective lineage analysis and determine a gastrula map of the origin of vascular endothelial cells from 6 to 27 hpf.

To construct single-cell resolution images of a zebrafish embryo, it is essential to capture light-sheet images from different viewing directions. The mounted embryo was illuminated first on the left side and then the right side, images were acquired *via* z-stacks from 0°, and the embryo was then rotated and the images were acquired from 180° (Fig. 1), thus generating four sets of imaging data at each time point (Fig. 2A, A’, and K). In this work, we captured three sets of image data from three embryos: embryo #1 (6–27 hpf), embryo #2 (6–22 hpf), and embryo #3 (7–27 hpf) (Supplementary Movies S1–S3). The raw data were ~5–7 terabytes (TB) in size (Supplementary Fig. S1). To fuse the data acquired by Zeiss Light Sheet Z.1, we first used the self-contained fusion module of Zen, the supporting software of Z.1. The Zen module is based on the image view settings on the microscope, but was not able to handle the two-view imaging (0° and 180°) of our samples and the imaging speed was too slow to keep up with the rapid development of zebrafish embryos. Moreover, the Zen module performed poorly on 4-view fusion (Supplementary Fig. S2). In addition, we found that for dual-illumination side-fusion, the only supportive strategies available were maximal, mean, and Fourier domain maximal, but all these measurements decreased the contrast of images. To address these questions, we created a software named AFEIO described in Materials and Methods (Fig. 2). We performed dual-side fusion (Fig. 2B and C), regional extraction (Fig. 2D and E), dual-view fusion (Fig. 2F–H), and alignment (Fig. 2I and J) in the Tg(*H2A.F/Z*:EGFP) channel. In addition, in the Tg(*kdrl*:mCherry) channel, we used a threshold based on a region where the signal intensity in the nuclei was strong (Supplementary Fig. S3A, A’, and B). Together, we were able to obtain time-lapse, shift-corrected, fused-in-whole panoramic image stacks of the embryos (Fig. 2K’; Supplementary Movies S4 and S5).

**Fig. 2.**
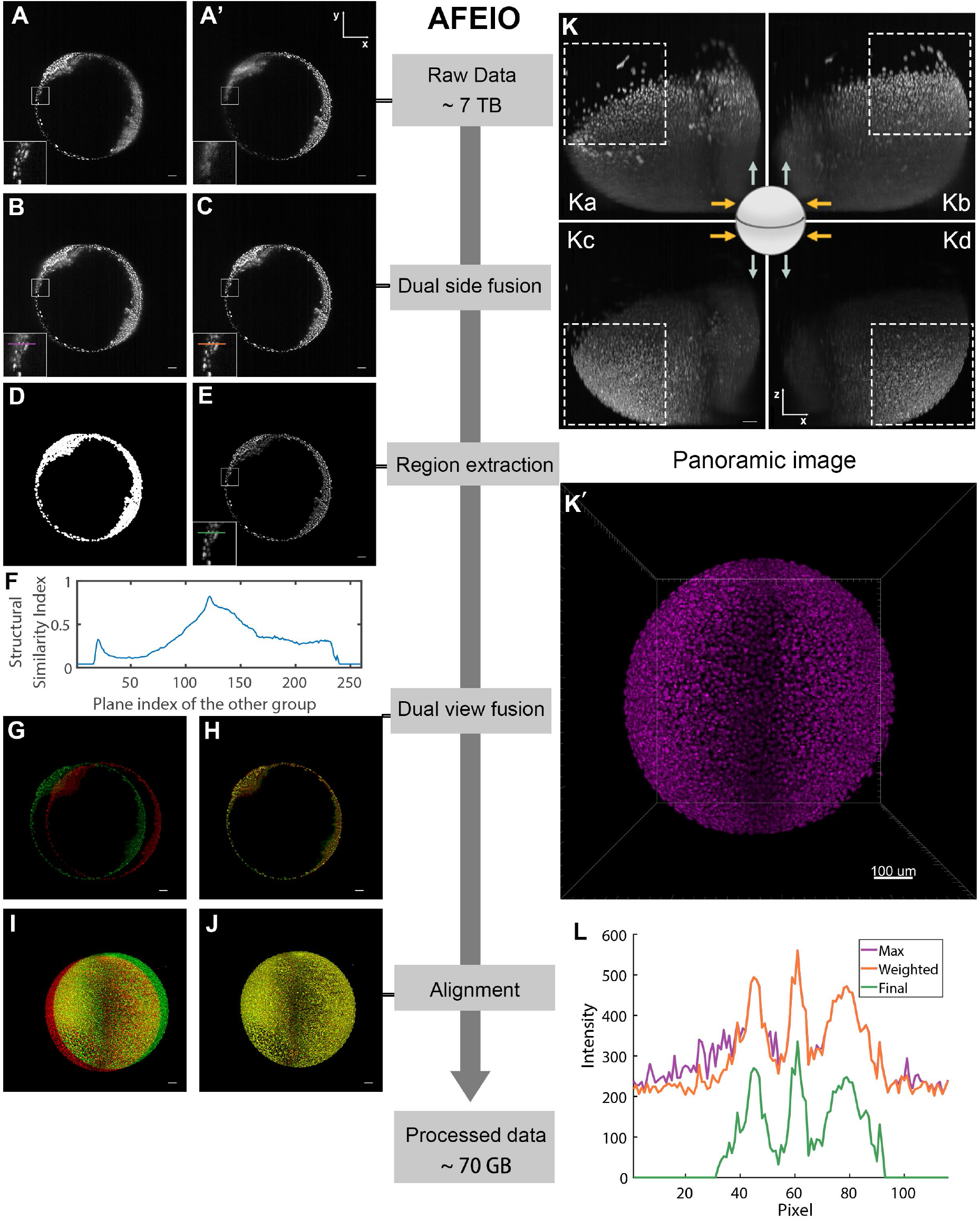
Single-cell resolution images of zebrafish embryos constructed from light-sheet data using AFEIO. The compressed, high-quality imaging data (~70 GB) are derived from the original raw data (~7 TB) using the AFEIO workflow, consisting of dual-side fusion, region extraction, dual-view fusion, and alignment. (A, A’) Original images of the same plane illuminated from the left (A) and right (A’). (B) Original image from the bi-directional illumination processed with ‘Max’ fusion. Enlarged square area refers to panel L for the intensity distribution along the purple line. (C) Original image from the bi-directional illumination processed with ‘Weighted’ fusion. Enlarged square area refers to panel L for the intensity distribution along the orange line. (D) The extraction mask, which only contains the pixels with information. (E) The image extracted from the ‘Weighted’ fusion image after masking. Enlarged square area refers to panel L for the intensity distribution along the green line. (F) Structural similarity of the selected plane from the 0° group and all planes from the 180° group. (G and H) A selected plane from the 0° group (red) and the most similar plane in the 180° group (green) (G) along with overlap after registration (H). (I and J) The shift between different time points (I) and overlap after registration (J). (K) Raw image projections of stacks from the top view with different illumination and imaging directions (orange arrows, illumination directions; green arrows, imaging directions). (Ka) Illumination from the left and detection at 0°; (Kb) illumination from the right and detection at 0°; (Kc) illumination from the left and detection at 180°; (Kd) illumination from the right and detection at 180°. (K’) Output of the fused imaging data (also shown in Movie S4). (L) Intensity distribution of maximal fusion, sigmoidal weighted fusion, and the final image; note that compared to the maximal fusion, the sigmoidal weighted fusion markedly improved the image contrast (Supplementary Fig. S3C). Scale bars in A-E, and G-K, 50 μm; K’, 100 μm.

### AFEIO software for time-lapse and dual-view image processing

It has been reported that the sigmoidal weighting function can be applied to dual-side fusion (de Medeiros et al., 2015). Compared with maximal fusion, we found that sigmoidal weighted fusion significantly improved the image contrast (Fig. 2L; Supplementary Fig. S3C). After background subtraction, an extraction mask was applied to the image, and only the regions where there were signals were retained (Fig. 2D-E). With advances in run-length encoding, the volume of data was significantly reduced in this step. In addition, Huisken and colleagues have already shown the feasibility of fusing light-sheet images taken from different directions (Huisken et al., 2004). Given that the interval between capturing images from two directions was very short compared with the duration of embryonic development in zebrafish, we regarded them as occurring simultaneously. Furthermore, there was sufficient overlap in the middle part, sharing the same spatial information, and this was used for static registration. Thus, we used the structural similarity index (Fig. 2F) to match the fused images corresponding to the same plane from the two directions (Zhou et al., 2004). Finally, to correct displacement at different time points, we first projected the main view and left view of a single embryo, then used the Fourier-Merlin transform to calculate the total displacement of the embryo in three dimensions between different time points, and corrected the displacement to achieve alignment (Fig. 2I and J). This image processing can be done on a personal computer equipped with a single i7-7700 CPU, 8 GB DDR3 memory, a 256 GB SSD, and a data-saving mobile hard disk. In addition to processing the datasets acquired from the Zeiss Z1 system, the AFEIO software and strategy were also able to process datasets from other light-sheet microscopes such as Luxendo Muvi-SPIM (Supplementary Movie S6). This processing strategy provided a high-contrast, low-redundancy dataset, reducing ~7 TB of raw data to ~70 GB for each of the three embryos (Supplementary Fig. S1) for further single-cell tracking of the origin of ECs during early development.

### Retrospective lineage tracing of the origin of vascular endothelial cells

The ~70 GB of high-quality data was then imported into “Imaris” software to calculate cell lineage relationships. We tracked the lineages of all EGFP^+^ cells and then filtered out EGFP^+^/mCherry^+^ ECs. The nuclei of the cells were labeled with Tg(*H2A.F/Z*:EGFP) and detected as spots by Imaris. The detailed locations of spots with tracks allowed us to follow each cell of a developing embryo (Supplementary Movie S7). Statistical analysis of embryo #1 showed the total number of cells doubled from 8176 (6 hpf) to 18698 (27 hpf) during early development (Supplementary Fig. S4). The number of cells at 18 hpf was 15905, close to that in a previous report (Keller et al., 2008), further validating the feasibility of our method. The total numbers of cells in embryo #2 were similar from 6 to 22 hpf (Supplementary Fig. S4). Therefore, these results suggest that our method is able to resolve *in toto* imaging of a zebrafish embryo consisting of >18,000 cells at single-cell resolution.

To trace the lineages of vascular ECs, we selected one EGFP^+^/mCherry^+^ double-positive nucleus near the eye at 27 hpf while simultaneously marking its sister nucleus (Fig. 3A). Retrospective fate-mapping from 27 to 6 hpf showed that these endothelial descendants in the head (Fig. 3A-D) migrated from the anterior lateral plate mesoderm (Fig. 3E), and originated from a single endothelial progenitor in the dorsal side of the shield embryo (Fig. 3F-H) (Supplementary Movie S8; yellow arrowhead). The lineage tree map showed that 27 ECs in the head at 27 hpf were traced back from 27 ECs at 23 hpf, 17 at 19 hpf, 8 at 15 hpf, 6 at 12 hpf, 5 at 10 hpf, 3 at 8 hpf, and a single dorsal endothelial progenitor at 6 hpf, suggesting dynamic cell division and death during early development (Fig. 3I). In parallel, we selected one EGFP^+^/mCherry^+^ double-positive nucleus in the posterior trunk at 27 hpf, and simultaneously marked its sister cells (Fig. 3J). Retrospective fate-mapping from 27 to 6 hpf showed that their endothelial descendants (Fig. 3J-M) migrated from the posterior lateral plate mesoderm (Fig. 3N), and traced back to one endothelial progenitor in the ventral side of the shield embryo (Fig. 3O-Q; yellow arrowhead; Supplementary Movie S9). The lineage tree map revealed that 6 selected ECs of the posterior trunk at 27 hpf were traced back from 2 ECs at 23 hpf, 1 at 19 hpf, 14 at 15 hpf, 13 at 12 hpf, 9 at 10 hpf, 2 at 8 hpf, and a single ventral endothelial progenitor at 6 hpf (Fig. 3R). To construct a comprehensive lineage map of the origins of all vascular ECs, we marked all the EGFP^+^/mCherry^+^ double-positive nuclei of ECs with different colors along the anterior-posterior axis from 18 to 22 hpf, and then performed lineage tracing back to the gastrula progenitors using Imaris. Taking the otic vesicle as the boundary between head and trunk, we marked vascular ECs of the head in red, and divided ECs of the trunk into two parts, the anterior trunk marked in blue and the posterior trunk marked in green (Fig. 4A).

**Fig. 3.**
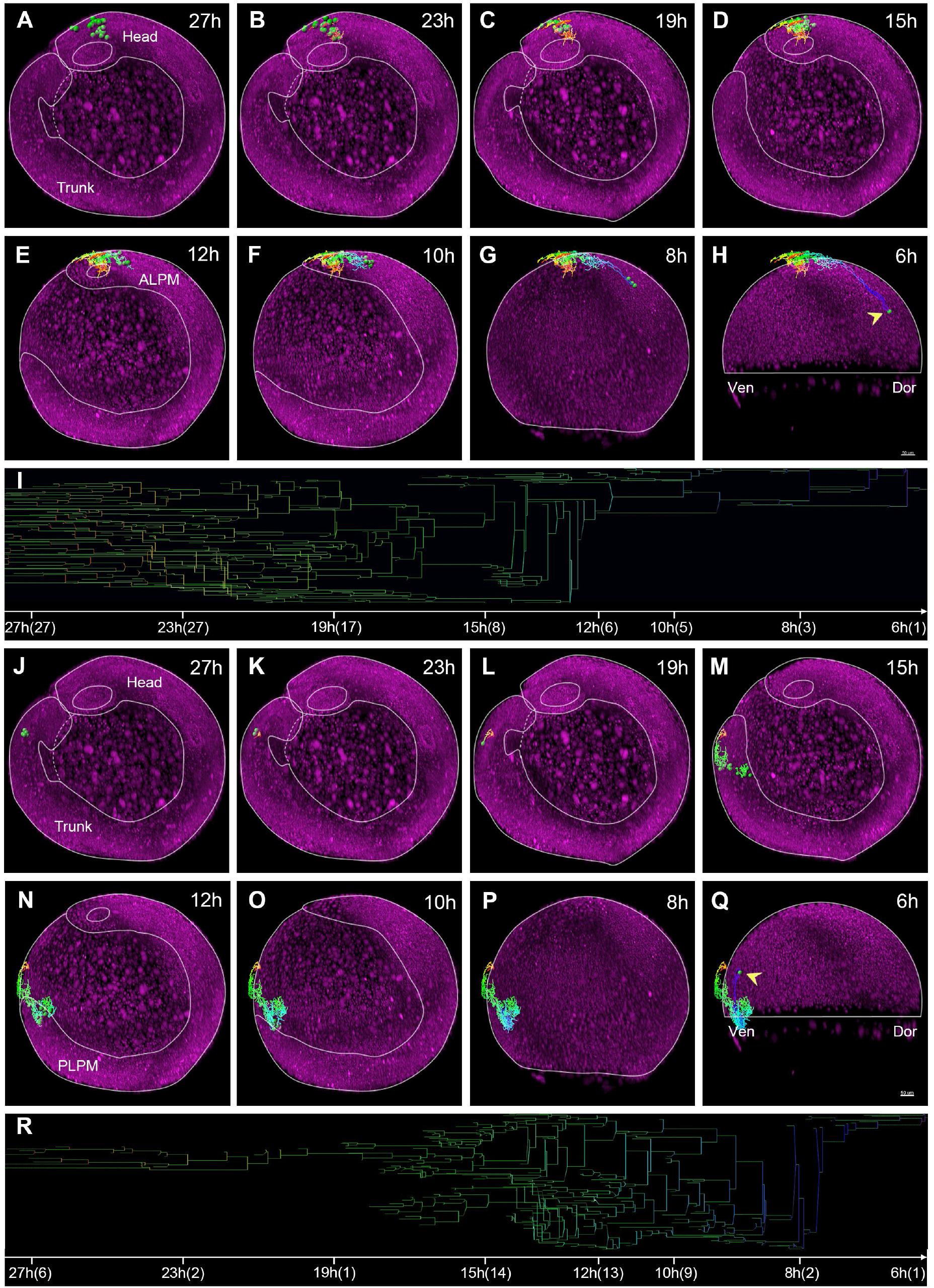
Retrospective cell-lineage tracking reveals the distinct gastrula origins of vascular endothelial cells in the trunk and head. (A-H) Representative images showing vascular endothelial cells of the head (green) at 27 hpf (A) retrospectively tracked back to a dorsal cell at 6 hpf (H, arrowhead), with endothelial cells in the head from 23 to 15 hpf (B-D), the anterior lateral plate mesoderm (ALPM) at 12 hpf (E), and the dorsal cells at 10 and 8 hpf (F, G). (I) Lineage tree map showing that the selected brain endothelial cells at 27 hpf are derived from a dorsal cell at 6 hpf. Along the time line, 27h(27), 27 ECs at 27 hpf; 23h(27), 27 ECs at 23 hpf, etc. (J-Q) Representative images showing vascular endothelial cells of the trunk at 27 hpf (J) retrospectively tracked back to a ventral cell at 6 hpf (Q), with endothelial cells in the trunk from 23 to 15 hpf (K-M), the posterior lateral plate mesoderm (PLPM) at 12 hpf (N), and the ventral cells at 10 and 8 hpf (O, P). (R) Lineage tree map showing that the selected trunk endothelial cells at 27 hpf derive from a single ventral cell at 6 hpf. Along the time line, 27h(6), 6 ECs at 27 hpf; 23h(2), 2 ECs at 23 hpf, etc. Yellow arrow points to the gastrula cells; multiple-colored lines showing the cell migration tracking; Scale bars, 50 μm.

**Fig. 4.**
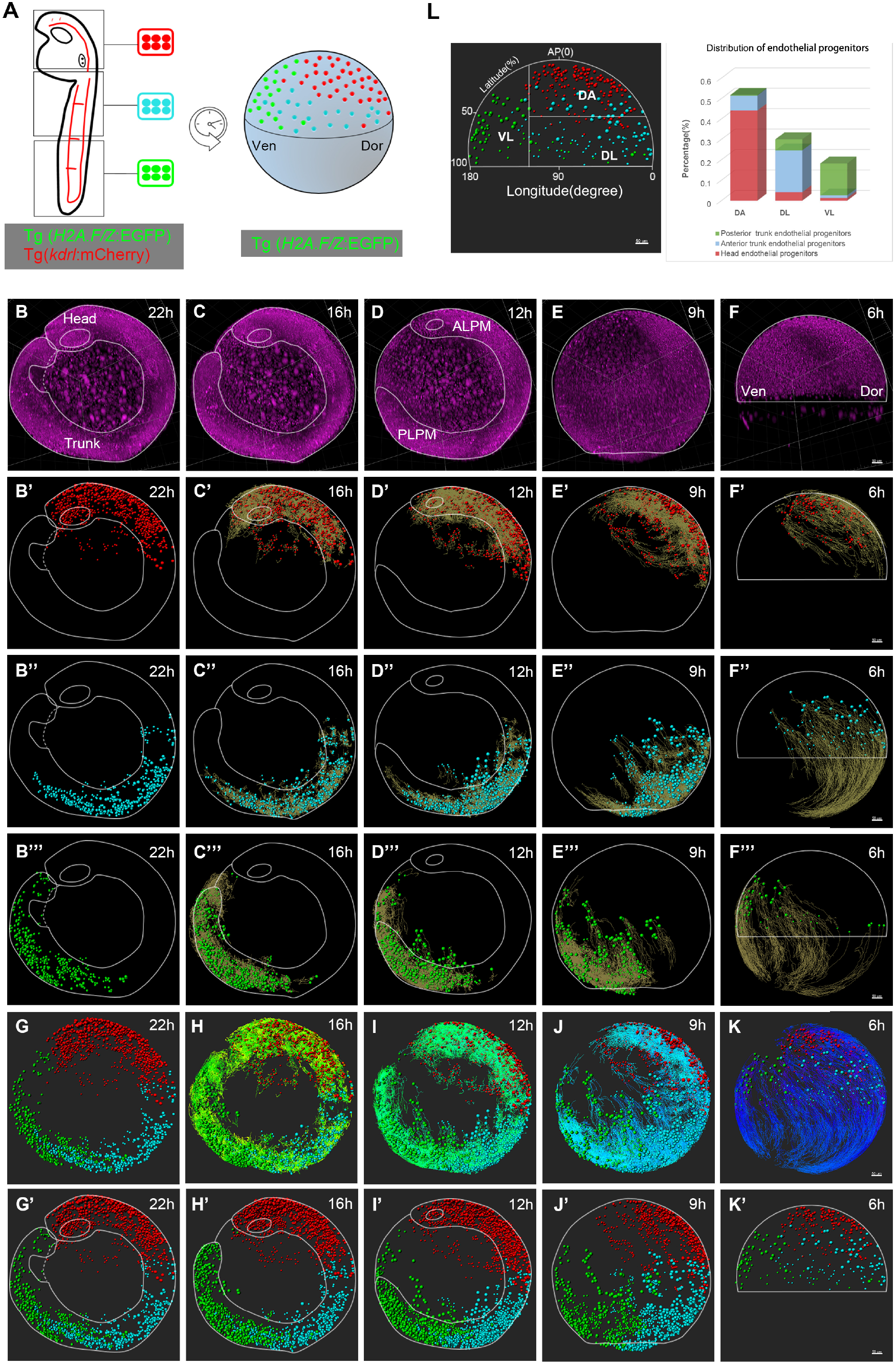
Retrospective cell-lineage tracing creates a comprehensive map of the origin of vascular endothelial cells along the dorsal-ventral (D-V) and anterior-posterior (A-P) axes of the gastrula. (A) Scheme for retrospective lineage tracing of the origin of vascular endothelial cells using Tg(*H2A.F/Z*:EGFP) to label the nuclei of all embryonic cells and Tg(*kdrl*:mCherry);Tg(*H2A.F/Z*:EGFP) to co-label the nuclei of all endothelial cells. All mCherry^+^ nuclei are classified with different colors: red in the head (the otic vesicle as the boundary between head and trunk), blue in the anterior trunk, and green in the posterior trunk. The gastrula progenitors at 6 hpf are retrospectively tracked from endothelial cells at different locations at 22hpf with Imaris software. Dor, dorsal; Ven, ventral. (B-F) Developmental morphology of embryo #1 at 22 hpf (B),16 hpf (C), 12 hpf (D), 9 hpf (E), and 6 hpf (F) (note the bend tail at 22 hpf) (ALPM, anterior lateral plate mesoderm; PLPM, posterior lateral plate mesoderm; Dor, dorsal; Ven, ventral). (B’-F’) Retrospective lineage tracing from the head endothelial cells at 22hpf (B’), 16 hpf (C’), 12 hpf (D’), and 9 hpf (E’) to the DA gastrula cells at 6 hpf (F’) (dynamic lineage tracking is shown in Supplementary Movie S10). (B’’-F’’) Retrospective lineage tracking from the anterior trunk endothelial cells at 22hpf (B’’), 16 hpf (C’’), 12 hpf (D’’), and 9 hpf (E’’) to the DL gastrula cells at 6 hpf (F’’) (dynamic lineage tracking in Supplementary Movie S11). (B’’’-F’’’) Retrospective lineage tracking from posterior trunk ECs at 22 hpf (B’’’), 16 hpf (C’’’), 12 hpf (D’’’), and 9 hpf (E’’’) to VL gastrula cells at 6 hpf (F’’’) (dynamic lineage tracking in Supplementary Movie S12). (G-K) Retrospective lineage tracking showing the distribution of the three clusters of ECs (red, blue, and green) with celllineage tracking lines (multiple-colored lines) from 22 hpf (G), 16 hpf (H), 12 hpf (I), and 9 hpf (J) to the gastrula progenitors at 6 hpf (K) (dynamic tracking in Supplementary Movie S14). (G’-K’) Retrospective lineage tracking showing the distribution of the three clusters of endothelial cells (red, blue, and green) without cell-lineage tracking lines from 22 hpf (G’), 16 hpf (H’), 12 hpf (I’), and 9 hpf (J’) to the gastrula progenitors at 6 hpf (K’). Scale bars, 50 μm. (L) Left panel: the gastrula divided into a dorsal–anterior (DA) region along the A–P axis (0–100% latitude), as well as dorsal–lateral (DL) and ventral–lateral (VL) regions along the D–V axis (0–180° longitude) (AP(0), animal pole as 0% latitude; scale bar, 50 μm). Right panel: quantitative analysis showing the percentage of the three regions (DA, DL, and VL) that contribute to ECs in the head (red), anterior trunk (blue), and posterior trunk (green). Note that the DA gastrula contributes to the head ECs, DL to the anterior trunk ECs, and VL to the posterior trunk ECs.

We then performed lineage analysis of the three groups of endothelial progenitors using retrospective fate mapping from 22 to 6 hpf, showing the developmental morphology at different stages (Fig. 4B-F; Supplementary Fig. S6; Supplementary Movie S1). The head vascular ECs from 22 hpf were traced back mainly to the anterior lateral plate mesoderm at 12 hpf (Fig. 4B’-D’; Supplementary Fig. S6A’-C’) and the DA gastrula progenitors near the animal pole at 6 hpf (Fig. 4D’-F’; Supplementary Movie S10; Supplementary Fig. S6C’-E’). The anterior trunk vascular ECs at 22 hpf were traced back to both the anterior and posterior lateral plate mesoderm at 12 hpf (Fig. 4B’’-D’’; Supplementary Fig. S6A’’-C’’) and the DL gastrula progenitors at 6 hpf (Fig. 4D’’-F’’; Supplementary MovieS11; Supplementary Fig. S6C’’-E’’). The posterior trunk vascular ECs at 22 hpf were traced back to the posterior lateral plate mesoderm at 12 hpf (Fig. 4B’’’-D’’’; Supplementary Fig. S6A’’’-C’’’) and the VL gastrula progenitors at 6 hpf (Fig. 4D’’’-F’’’; Supplementary Movie S12; Supplementary Fig. S6C’’’-E’’’). In addition, the vascular EC migration and division tracks were shorter between 22 and 12 hpf than those between 12 and 6 hpf, suggesting rapid derivation, division, and migration of ECs along the body axes from 6 to 12 hpf (Supplementary Movie S13, yellow lines). The combined lineage maps of all three groups of endothelial progenitors were projected onto representative images with lineage tracing lines (Fig. 4G-K; Supplementary Fig. S6F-J) or without them (Fig. 4G’-K’; Supplementary Fig. S6F’-J’) as well as movies (Supplementary Movies S14 and S15). The lineage tree maps of all ECs from embryo #1 were constructed in the scalable vector graphics (SVG) format and were included as a supplementary file. Quantitative analysis of two sets of data from embryos #1 and #2 revealed that vascular ECs derived from broadly-distributed endothelial progenitors of the shield embryo (Fig. 4L, left panel from embryo #1; Supplementary Fig S6J’ from embryo #2). The dorsal-anterior (DA) gastrula contributed to the formation of ECs mainly in the head and seldom in the anterior trunk; the dorsal-lateral (DL) gastrula gave rise to ECs mainly in the anterior trunk and seldom in the posterior trunk and head; and the ventral-lateral (VL) gastrula contributed to ECs mainly in the posterior trunk (Fig. 4L; right panel from embryo #1). Therefore, in contrast to the previous report that vascular ECs are primarily derived from the VL gastrula cells (Gore et al., 2012; Vogeli et al., 2006), our results support the conclusion that endothelial progenitors are distributed throughout the gastrula along the dorsal-ventral and anterior-posterior axes (Fig. 4K and K’; Supplementary Fig. S6J and J’); the DA gastrula cells give rise to most of the ECs in the head while the DL and VL gastrula cells contribute to most of the ECs in the trunk and tail.

Importantly, this digital cell-lineage mapping enabled the visualization of all cell divisions, migration, and differentiation at single-cell resolution. We randomly selected a single cell near the animal pole of the shield embryo (Supplementary Fig. S7A), which started to rapidly divide and migrate upwards to the animal pole and then migrated to the right (Supplementary Fig. S7B). At ~9 hpf, its descendants began to migrate back (Supplementary Fig. S7C and D) and further migrated to the upper left of the starting position (Supplementary Fig. S7E and F), finally reaching the dorsal head between the eyes (Supplementary Fig. S7G and H; Supplementary Movie S16). The lineage tree map showed dynamic cell division, migration, and death from 6 to 27 hpf (Supplementary Fig. S7I). Thus, if the cells of interest in embryonic organs are marked with a fluorescent reporter, the method described here enables tracing them back to their gastrula origin in early zebrafish embryos.

### Prospective lineage tracing the origin of vascular ECs using the Kaede reporter

To confirm the distinct gastrula origin of vascular ECs between the head and trunk/tail, we used the photo-convertible fluorescent protein Kaede to analyze prospective cell lineages as previously reported (Ando et al., 2002). We first established a Tg(*kdrl*:kaede) transgenic zebrafish line in which *kaede* was driven by the *kdrl* promoter. We generated several independent Tg(*kdrl*:kaede) transgenic lines from different F0 founders and all of them had similar kaede expression patterns, with quite strong expression at 6 hpf and later in blood vessels. Whole-mount *in situ* hybridization and semi-quantitative PCR revealed that *kdrl* was expressed during gastrulation (4 hpf at the earliest) and several VEGF (vascular endothelial growth factor) ligands were also expressed in early embryos (Supplementary Fig. S8), which is consistent with previous reports in zebrafish (Bussmann et al., 2007) and in mice (Ishitobi et al., 2011; Yamaguchi et al., 1993). In addition, kaede expression in the anterior lateral plate mesoderm in Tg(*kdrl*:kaede) transgenic embryos partly overlapped with that in Tg(*scl-α:*dsRed) embryos (Supplementary Fig. S9) (Zhen et al., 2013). These data suggested that Tg(*kdrl*:kaede) recapitulates the endogenous *kdrl* expression pattern. Upon photoactivation by a 405-nm laser, a small cluster of Kaede^+^ cells in the dorsal gastrula at 6 hpf were converted from fluorescent green to red (Supplementary Fig. S10A). The development of these red cells and the other green Kaede^+^ cells were followed by confocal microscopy at 12, 20, and 28 hpf (Supplementary Fig. S10B-D). We noted that the red cells divided and migrated to the anterior lateral plate mesoderm at 12 hpf (Supplementary Fig. S10B), continued to migrate to the head region, and eventually formed vascular endothelium there (Supplementary Fig. S10C and D). In contrast, when a small group of ventral gastrula cells were photo-activated from Kaede^+^ fluorescent green to red (Supplementary Fig. S10E), these red cells divided and migrated to the posterior lateral plate mesoderm at 12 hpf (Supplementary Fig. S10F), then continued to migrate, and eventually formed vascular ECs in the trunk and tail (Supplementary Fig. S10G and H). Therefore, these data further support the conclusion that vascular ECs are derived from different gastrula cells along the body axis, the fate of VL gastrula cells being ECs in the trunk/tail and that of DA gastrula cells being ECs in the head.

## Discussion

We have developed an effective workflow for tracing cell lineages in zebrafish using a Zeiss Z.1 LSFM, Imaris software, and AFEIO software. In the multi-view registration fusion field, many investigators have already reported well-developed software and/or protocols. A comprehensive multi-view image fusion pipeline was first reported in 2007 (Swoger et al., 2007), and a bead-based registration software was later reported (Preibisch et al., 2010). The IsoView image processing software provides a high processing speed, isotropic resolution, and a high compression rate (Amat et al., 2015; Chhetri et al., 2015). The efficient Bayesian multiview deconvolution method significantly accelerated the image processing speed (Guo et al., 2019; Preibisch et al., 2014; Wu et al., 2013). The Massive Multi-view Tracker (MaMuT) software enables the reconstruction of cell lineages at single-cell resolution (Wolff et al., 2018). Our AFEIO software built upon the existing imaging registration and fusion software and incorporated additional functionalities to better suit applications for early zebrafish embryos. Therefore, the AFEIO software enables to construct a high-contrast, low-redundancy, fused-in-whole, and time-lapse embryo datasets, thus revealing the very first comprehensive digital map of the origin, division, and migration of ECs in zebrafish embryos from 6 to 27 hpf.

This digital map elucidates the contribution of the DA gastrula to ECs in the head while the VL and DL gastrula to ECs in the trunk and tail as summarized in Figure 5. Our working model suggests that vascular ECs are derived from broad areas of gastrula cells and are not limited to the VL gastrula cells (Vogeli et al., 2006). This is probably due to the light-sheet technology development, which allowed us to map all the gastrula regions with much higher numbers of tracked cells (n>200) while the previous works primarily mapped the fates of the ventral-lateral region of the gastrula. In addition, vascular endothelial progenitors were distributed along the anterior-posterior axis at 12 hpf, suggesting that they were likely not restricted to the anterior and posterior lateral plate mesoderm that warrants an investigation in the future. The distinct origin of cerebral and trunk endothelial cells revealed by this work is consistent with that the development of cerebral and trunk vessels is regulated by distinct signaling pathways as previously reported (Stenman et al., 2008). Regarding the major celllineage tracing methods, dye injection can only trace a few cell-divisions (Kimmel et al., 1990); tissue-specific transgenic reporters or genetic recombination methods are heavily dependent on the availability of the promoters/enhancers required to label both progenitors and descendants (Kretzschmar and Watt, 2012; Spanjaard and Junker, 2017); and retroviral- or genome editing-based barcodes are able to trace long-term cell lineage relationships but do not capture spatial information (Kalhor et al., 2017; McKenna et al., 2016; Spanjaard and Junker, 2017). Our method resolves both long-term and spatial light-sheet images by only using a transgenic reporter that labels the descendants/differentiated cells of interest, such as the *kdrl*:mCherry-labelled ECs in this work. Together, this work not only deciphers the distinct gastrula origins of cerebral and trunk ECs in zebrafish (Schoenebeck et al., 2007) but also our method has the potential for broad applications in decoding the origins of organs in zebrafish and other model organisms.

**Fig. 5.**
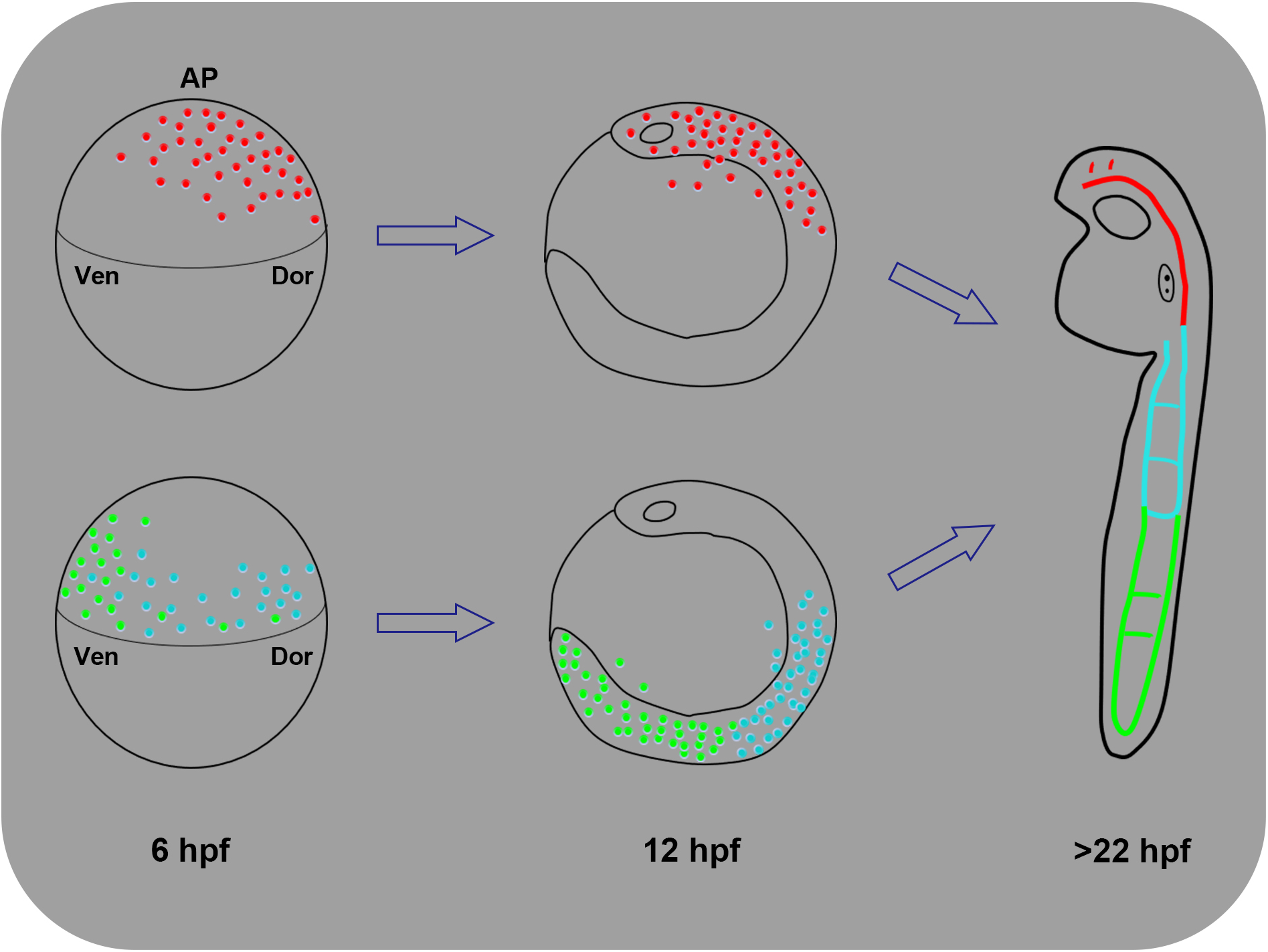
A new lineage-map model of the gastrula origin of vascular ECs in zebrafish. Based on this work, we propose that the dorsal-anterior gastrula cells (red) contribute to brain ECs, the dorsal-lateral gastrula cells (blue) to anterior trunk ECs, and the ventral-lateral gastrula cells (green) to posterior trunk ECs. This new model suggests that vascular ECs are derived from broad areas of the gastrula and are not limited to the ventral-lateral gastrula cells. AP, animal pole; Dor, dorsal; Ven, ventral.

## Materials and Methods

### Zebrafish lines and maintenance

Zebrafish were raised and handled in accordance with the animal protocol *IMM-XiongJW-3* approved by the Peking University Institutional Animal Care and Use Committee accredited by the Association for Assessment and Accreditation of Laboratory Animal Care International. The Tg(*kdrl*:kaede) transgenic line was created to express Kaede under the control of the vascular endothelium-specific *kdrl* promoter (Choi et al., 2007) using the Tol2 transposon elements (Kawakami et al., 2000), and the Tg(*kdrl*:mCherry) line (Xia et al., 2013) was from Dr. Bo Zhang (School of Life Science, Peking University, Beijing, China). The Tg(*H2A.F/Z*:EGFP) line was kindly provided by Dr. Qiang Wang (Institute of Zoology, Chinese Academy of Sciences, Beijing, China) (Pauls et al., 2001).

### Preparation of fluorinated ethylene propylene (FEP) tubes and wire plungers

FEP tubes (S1815-04; Bola, Grünsfeld, Germany) were rinsed sequentially with 1 M NaOH and 0.5 M NaOH, ultrasonicated, rinsed in double-distilled water and 70% ethanol, and ultrasonicated again (Kaufmann et al., 2012). Finally, the tubes were cut and stored in 50-ml tubes. After 3 rinses in double-distilled water, each tube was coated with 3.0% methylcellulose before use. The wire plungers (701998; BRAND) were cleaned with 70% ethanol and stored in an autoclaved container.

### Optimized mounting of zebrafish embryos

Tg(*H2A.F/Z:EGFP*)^Tg/+^; Tg(*kdrl*:mCherry)^Tg/+^ double transgenic zebrafish embryos were maintained at 28.5°C. At 4 hpf, the embryos were carefully dechorionated with pronase (11458643001; Roche), followed by 5 washes with E3 medium. Then, each embryo was carefully transferred into 0.2% ultrapure low melting-point agarose (16520-050; Invitrogen) without tricaine. The embryo was then drawn into an FEP tube using a 1-ml syringe with an 18G blunt needle. We used a razor-blade to cut the tube from the needle, and retained the segment of tube containing the embryo (~1 cm long). After solidification of the agarose, the tube was fixed to a fine wire plunger with glue and Parafilm (PM-996; Bemis, Oshkosh, WI). Before imaging, the extra FEP was cut from both ends to ensure that the tube was filled with agarose and to avoid the formation of bubbles. The tube was cut as short as possible to reduce fluctuations during rotation and guarantee E3 infiltration. The ends of the wire plunger without FEP were wrapped with Parafilm (PM-996; Bemis) and fixed in a capillary (701908; BRAND). Finally, the capillary was fixed in a sample holder for subsequent imaging (Fig. 1).

### Long-term live imaging by light-sheet fluorescence microscopy

The images were acquired with a Zeiss Z.1 LSFM, which recorded one-view images in 3-μm Z steps and 260 images per time point, thus taking 30–35 s, which meant that the temporal resolution permitted no more than two viewing directions. We used two 5× illumination objectives, a 10× imaging objective, and 488-nm and 561-nm lasers. First, we adjusted the longest diameter of the wire plunger in the FEP tube to be perpendicular to the objectives, thus avoiding contact with them during sample rotation and imaging. Second, we adjusted the sheet position and image parameters, and set the image range. The image size was 1920 ×1920 pixels. The step size in the Z axis was 3 μm and 260 images were obtained in one view; the initial view was set as group 1 (0°), then the sample was rotated 180° and set as group 2 (180°). The temperature during imaging was maintained at 28.5°C. From 6 to 10 hpf, the time interval between views was 90 s and the camera only detected the EGFP signals of the double Tg (H2A.F/Z:EGFP); Tg(*kdrl*:mCherry) transgenic embryos. At ~10 hpf, the mCherry signal appeared, so we turned on the 561-nm laser and the camera simultaneously acquired both EGFP and mCherry signals at 150-s intervals. After 12 hpf, tricaine (200 mg/L) was applied to anesthetize the embryos. The images of embryos from 6 to 27 hpf were acquired and initially stored in a computer workstation for the light-sheet microscope.

### Lineage tracing with Imaris software

The above processed data were imported into Imaris. First, we performed lineage tracking of all embryonic cells with the EGFP channel. Second, we filtered the mCherry channel colocalization with the EGFP channel to mark ECs and their progenitors. Then we selected a single or several endothelial progenitors of interest to present the track and fate of these cells, and we labeled ECs at different positions with different colors from 15 to 27 hpf. Using the otic vesicle as the boundary between head and trunk, we marked the vascular ECs of the head in red and divided the ECs of the trunk into two parts, anterior marked in blue and posterior marked in green. When this labeling was completed, ECs at the different locations were clearly distinguishable. In the data tab, the numbers of both total embryonic cells and ECs at different time points were exported in Excel format.

### Qualitative analysis of endothelial cells with MATLAB

To qualitatively analyze the numbers of endothelial progenitors at different locations in the dorsal and ventral regions of the gastrula at 6 hpf, we used a MATLAB script. First, we set the size of a single cell as 1 pixel in Imaris and exported the images. Second, we separated the different colors in the original images. The cells were separated into three colors: red, green, and blue, whose RGB values were [255,0,0], [0,255,0], and [0,255,255], respectively. We used R = 255 to separate the red cells, B = 255 to separate the blue cells, and counted the remaining cells as green. Finally, we counted the numbers of single-color cells. By binarizing the images, we then considered the pixel gray level as 0 where there were no cells and as 1 where cells were present. By adding up the gray levels of the images, we obtained the numbers of cells of each color in the dorsal and ventral regions.

### UV-induced photoconversion of Kaede for fate mapping of vascular endothelial cells

Kaede is fluorescent green but can be photo-converted to fluorescent red by violet or UV light (Ando et al., 2002). Tg(*kdrl*:kaede) transgenic embryos with strong fluorescent signals were mounted in 0.35% low-melting-point agarose (Sigma). These embryos were photo-activated using a Nikon A1R microscope (Nikon, Tokyo, Japan), and captured images were processed with the 3D projection feature of NIS-Elements software. Briefly, at 6 hpf, a group of Kaede^+^ cells were activated with a 405-nm laser until their fluorescent green signals were nearly absent. These embryos were then imaged *via* both the 488-nm and 561-nm channels at 12, 20, and 28 hpf while anesthetized by tricaine.

## Acknowledgments

The authors thank Drs IC Bruce and Jiaye He for critical comments and reading the manuscript, and Drs Liu Yiqun and Fu Liqin for assistance with LSFM at the National Center for Protein Sciences at Peking University in Beijing, China. This work was supported by grants from the National Natural Science Foundation of China (31730061, 31327901, 31430059, 81470399, 81870198, and 31821091) and the National Key R&D Program of China (2018YFA0800500, 2019YFA0801600).

## Author Contributions

MP and LLB performed experiments, analyzed data, and wrote the manuscript; WJZ, XW, YB, CX, JYZ, and JYL helped with experiments and analyzed data; WG, ZF, LC, JZ, and HC helped with light-sheet fluorescence microscopy imaging, data processing, and data analysis; and XZ and JWX conceived and designed this work, analyzed data, and wrote the manuscript.

## Availability of Data and Materials

All data and materials are available in the manuscript, the Supplementary Information, and below websites: https://disk.pku.edu.cn/#/link/4ED92786A171450C943A370BBC545007 Or https://www.youtube.com/playlist?list=PLK9xesWj3KRk3rt0nshxQOCokbqCSI2_P AFEIO software and its user instruction can also be downloaded on the Github website: https://github.com/Manearth/AFEIO

Further information and requests for resources and reagents should be directed to and will be fulfilled by corresponding author.

## Supplementary Information

### List of Supplementary Information

1. Supplementary Figs. S1 to S10
2. Supplementary Movies S1 to S16, AFEIO software and its user instruction, and a file of cell-lineage maps for all endothelial cells in the SVG format can be downloaded: https://disk.pku.edu.cn/#/link/4ED92786A171450C943A370BBC545007 Or https://www.youtube.com/playlist?list=PLK9xesWj3KRk3rt0nshxQOCokbqCSI2_P
3. AFEIO software and its user instruction can also be downloaded on the Github website: https://github.com/Manearth/AFEIO

### Supplementary Figures (Figs S1-10)

**Supplementary Fig. S1.**
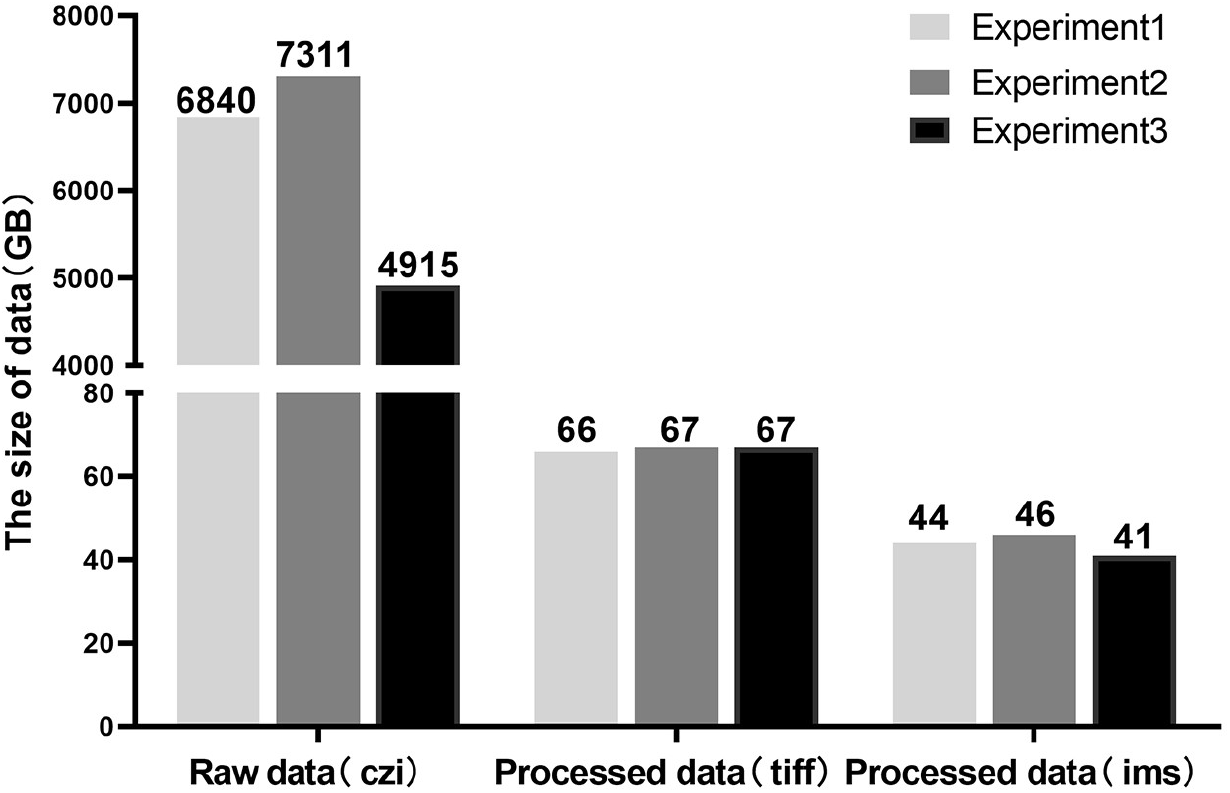
The size of light-sheet imaging data decreases ~100 fold after AFEIO processing. The datasets collected from three independent imaging experiments, showing that the raw data in Carl Zeiss image (czi) format is ~5–7 TB, and the data processed by AFEIO in tiff format is <70 GB or in Imaris (ims) format is <50 GB.

**Supplementary Fig. S2.**
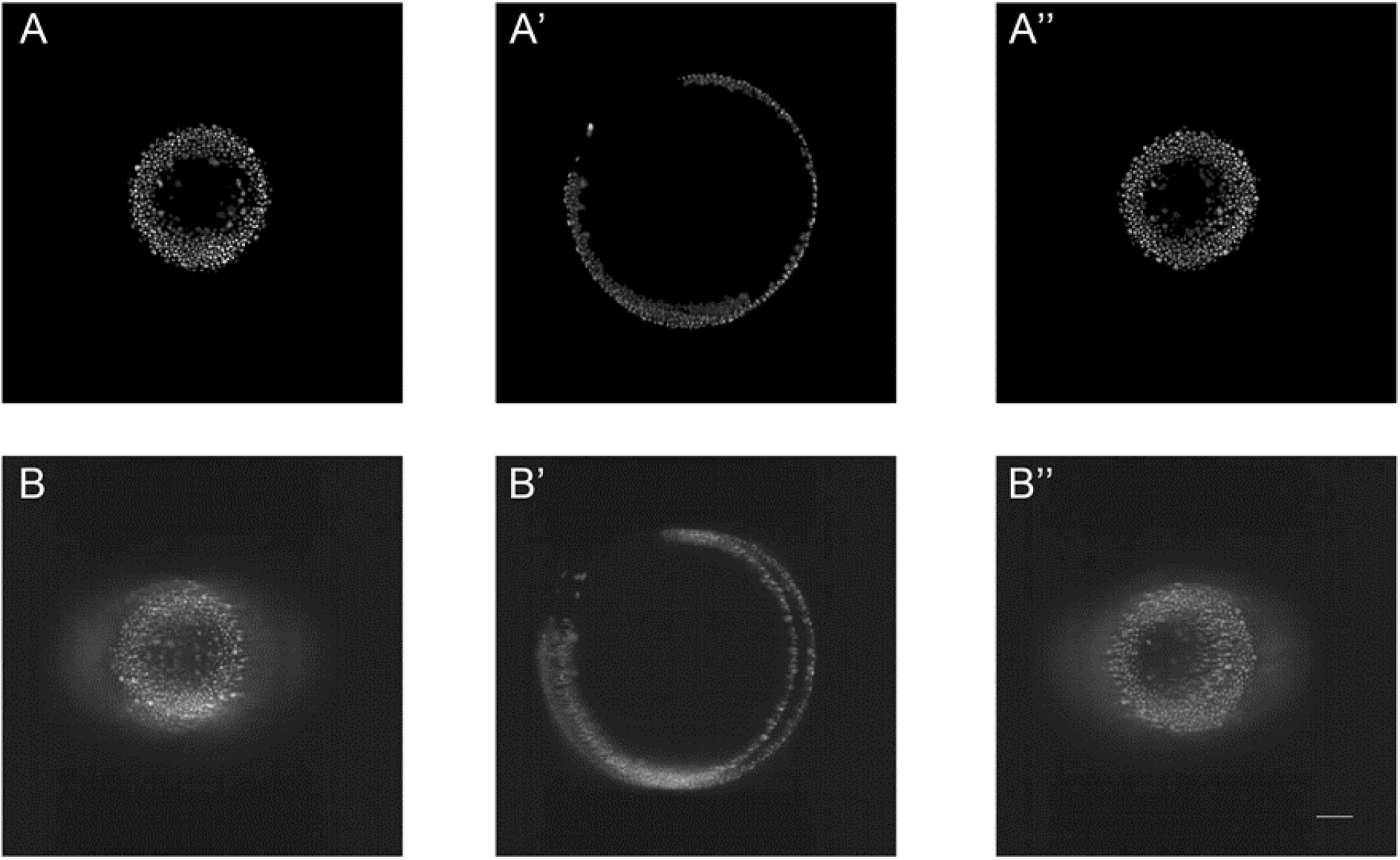
The image fusion by AFEIO (two-view imaging) results in higher resolution images than by Zen (four-view imaging). Three different planes were either fused by AFEIO (A, A’, A’’) or by Zen (Zeiss) (B, B’, B’’); note that AFEIO, but not Zen, generated high-quality image fusion.

**Supplementary Fig. S3.**
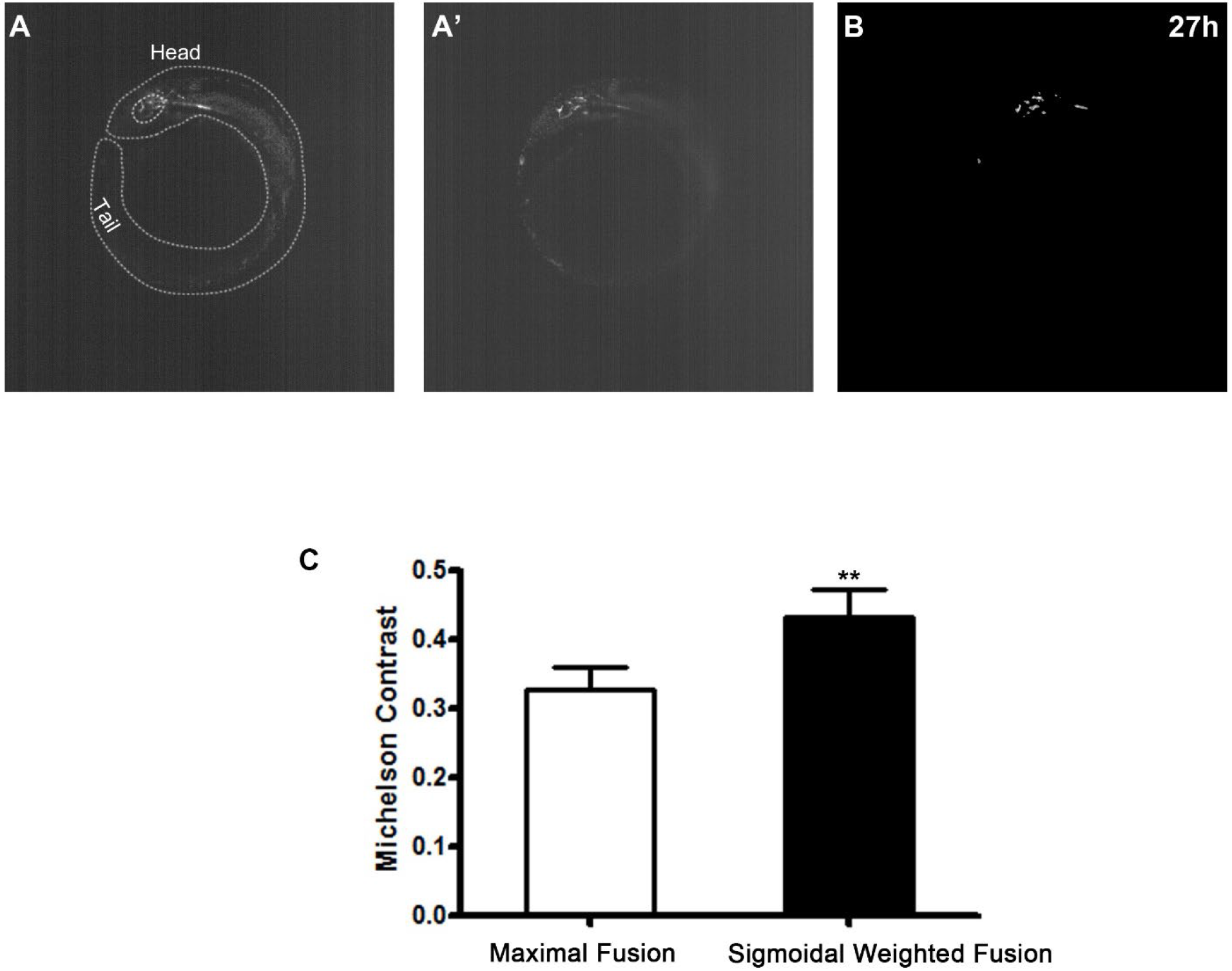
Image processing by AFEIO. (A, A’) Original Tg(*kdrl*:mCherry) images of embryo at 27 hpf, illuminated from the right (A) and left (A’). (B) The processed image after fusion and thresholding. (C) Michelson contrast with maximal fusion and sigmoidal weighted fusion; note that the latter significantly improves the contrast (p = 0.0094; n = 5).

**Supplementary Fig. S4.**
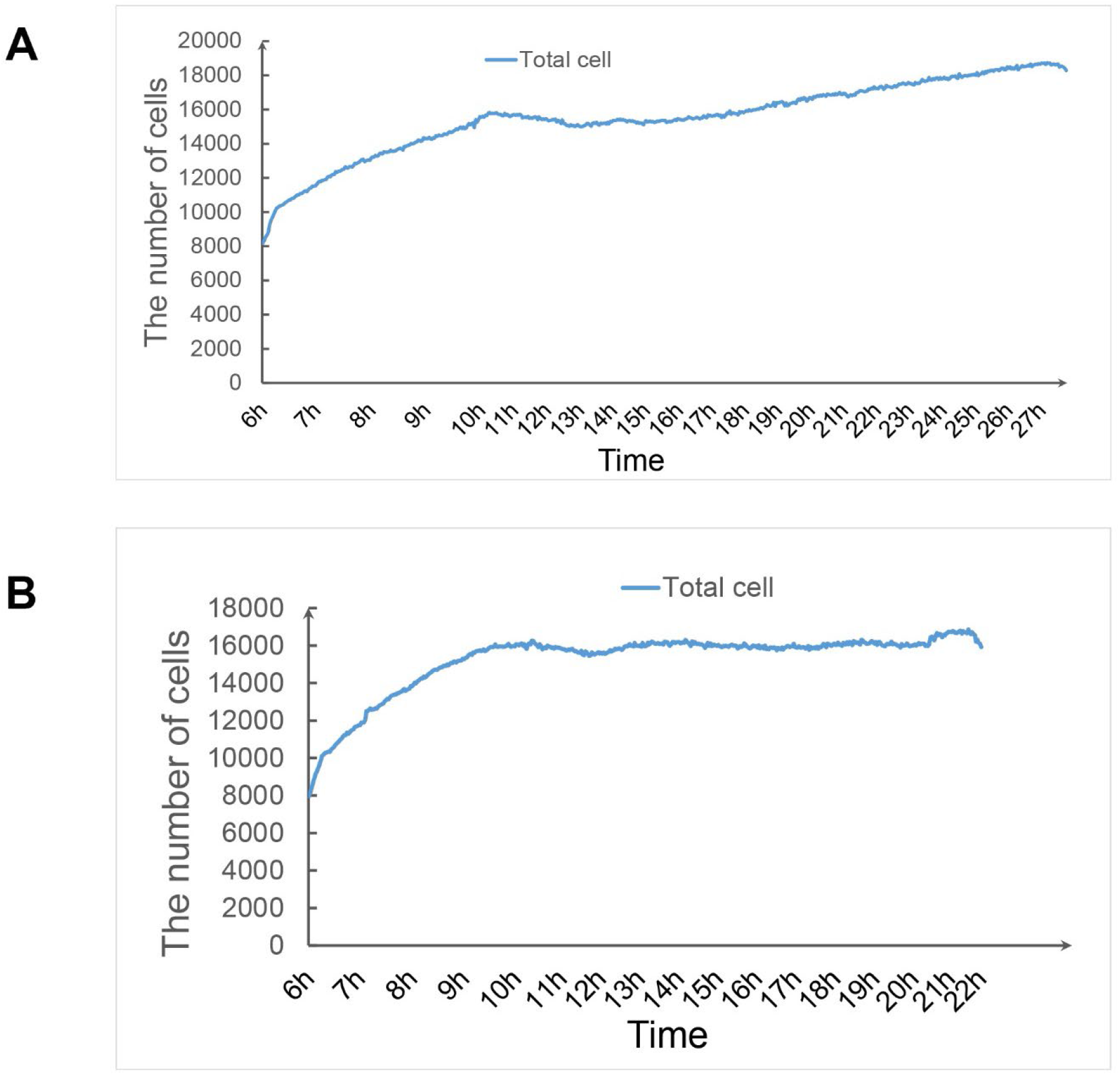
Dynamic changes in the total number of embryonic cells during early embryogenesis. The total numbers of cells were ~8000 at 6 hpf, increased quickly until 10 hpf, then decreased from 10 to 12 hpf, probably due to apoptosis, and later increased slowly until the end of imaging in embryo #1 (A) and embryo #2 (B).

**Supplementary Fig. S5.**
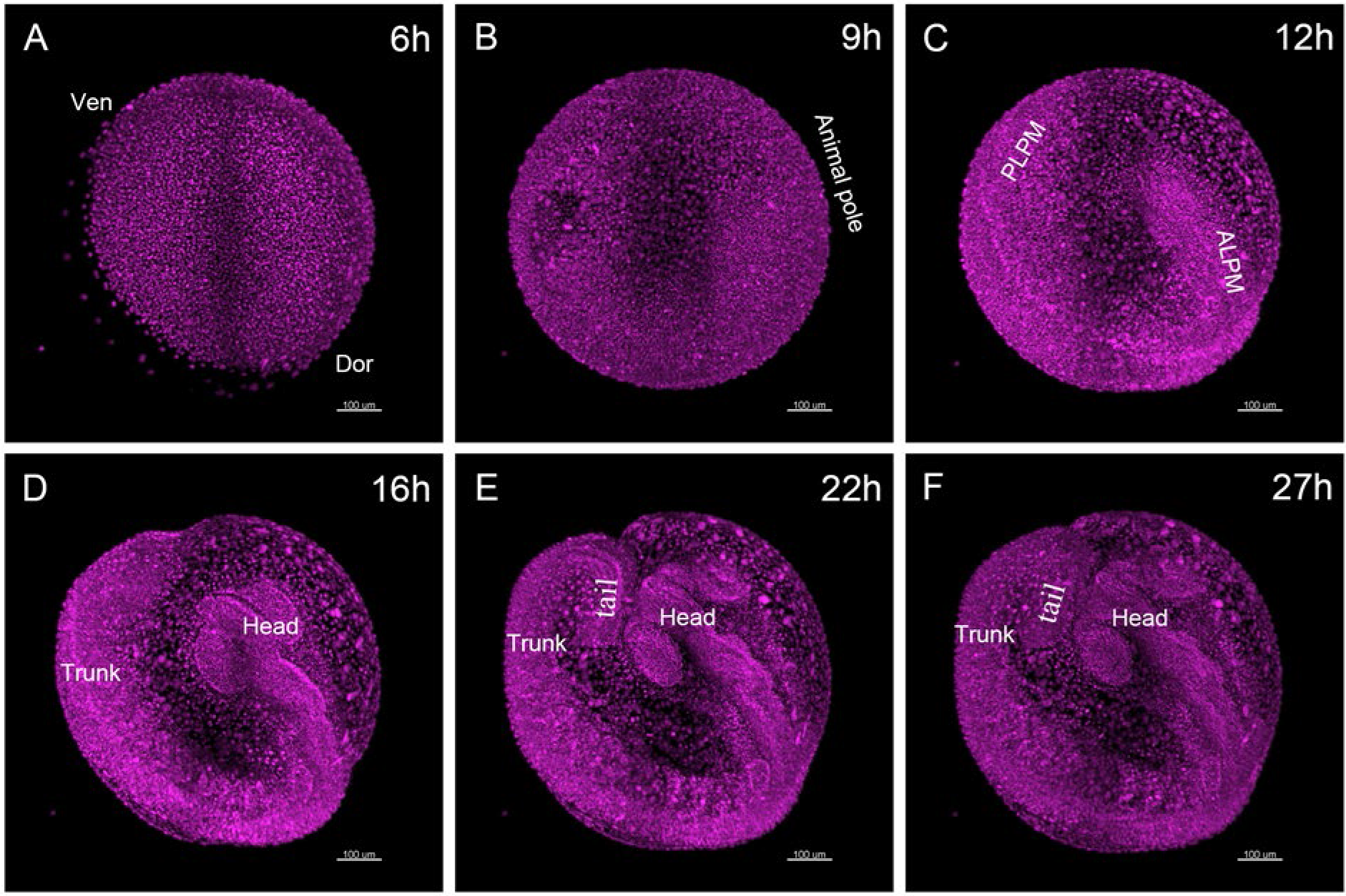
Developmental morphogenesis of embryo #1 from 6 to 27 hpf (also shown in Supplementary Movie S1). (A–F) Side view of the embryo at 6 hpf (A), 9 hpf (B), 12 hpf (C), 16 hpf (D), 22 hpf (E) and 27 hpf (F); note that the embryo develops normally but is not able to stretch out, with the growing tail bent (E–F) because it is embedded in FEP at an inner diameter of only 0.8 mm. Dor, dorsal; Ven, ventral; ALPM, anterior lateral plate mesoderm; PLPM, posterior lateral plate mesoderm. Scale bars, 100 μm.

**Supplementary Fig. S6.**
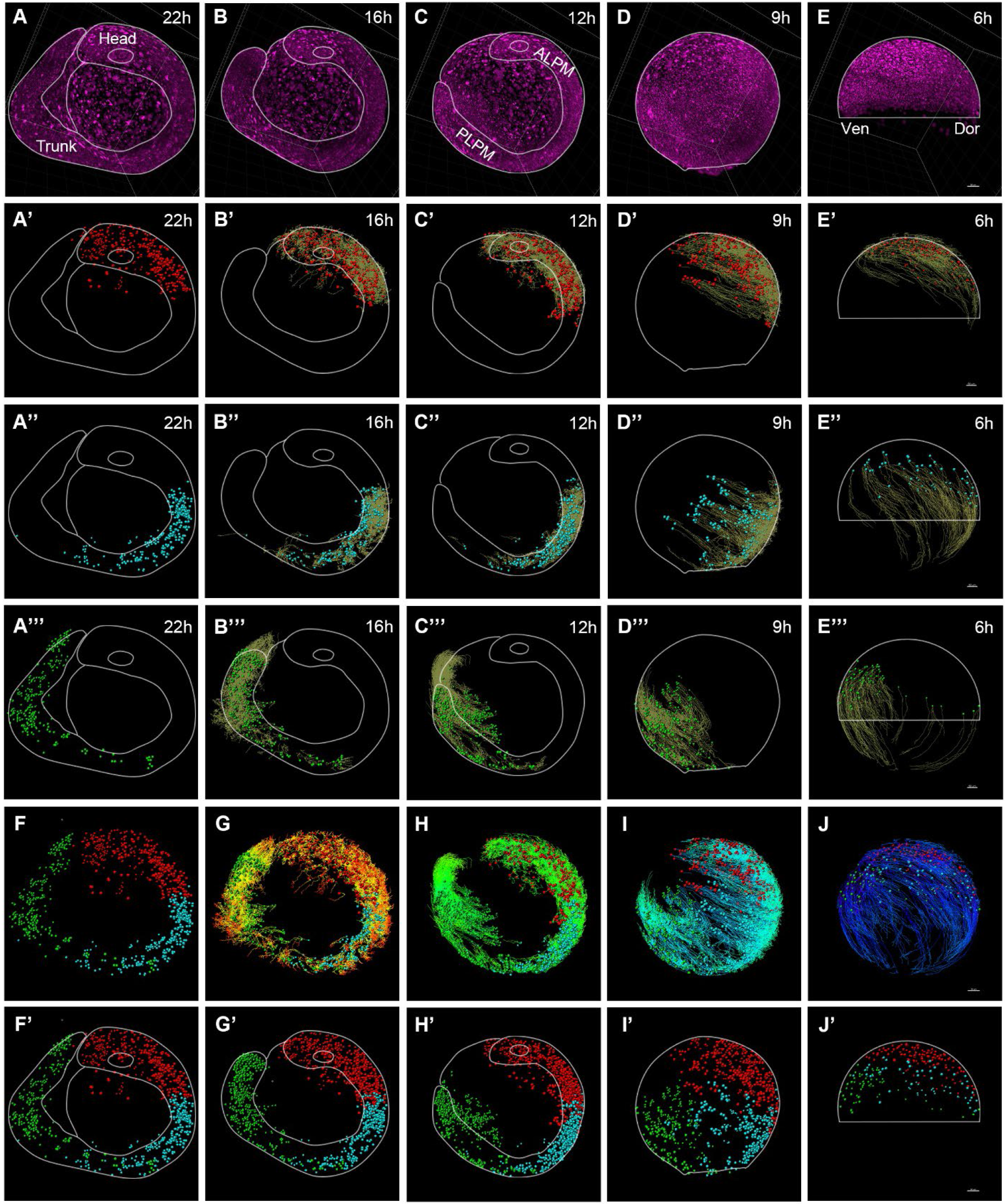
Retrospective cell-lineage tracing creates a comprehensive map of the origin of vascular endothelial cells along the dorsal–ventral and anterior–posterior axes of the gastrula in embryo #2. (A–E) Developmental morphogenesis of embryo #2 at 22 hpf (A),16 hpf (B), 12 hpf (C), 9 hpf (D), and 6 hpf (E). (A’–E’) Retrospective lineage tracing of head endothelial cells at 22 hpf (A’), 16 hpf (B’), 12 hpf (C’), and 9 hpf (D’) to the gastrula progenitors at 6 hpf (E’). (A’’–E’’) Retrospective lineage tracing of anterior trunk endothelial cells at 22 hpf (A’’), 16 hpf (B’’), 12 hpf (C’’), and 9 hpf (D’’) to gastrula progenitors at 6 hpf (E’’). (A’’’–E’’’) Retrospective lineage tracing of posterior trunk endothelial cells at 22 hpf (A’’’), 16 hpf (B’’’), 12 hpf (C’’’), and 9 hpf (D’’’) to the gastrula progenitors at 6 hpf (E’’’). (F–J) Retrospective lineage tracing showing the distribution of the three clusters of endothelial cells (red, blue, and green) with lineage tracking lines (multiple-colored lines) from 22 hpf (F), 16 hpf (G), 12 hpf (H), and 9 hpf (I) to gastrula progenitors at 6 hpf (J). (F’–J’) Retrospective lineage tracing showing the distribution of the three clusters of endothelial cells (red, blue, and green) without lineage tracking lines from 22 hpf (F’), 16 hpf (G’), 12 hpf (H’), and 9 hpf (I’) to the gastrula progenitors at 6 hpf (J’). Scale bars, 50 μm.

**Supplementary Fig. S7.**
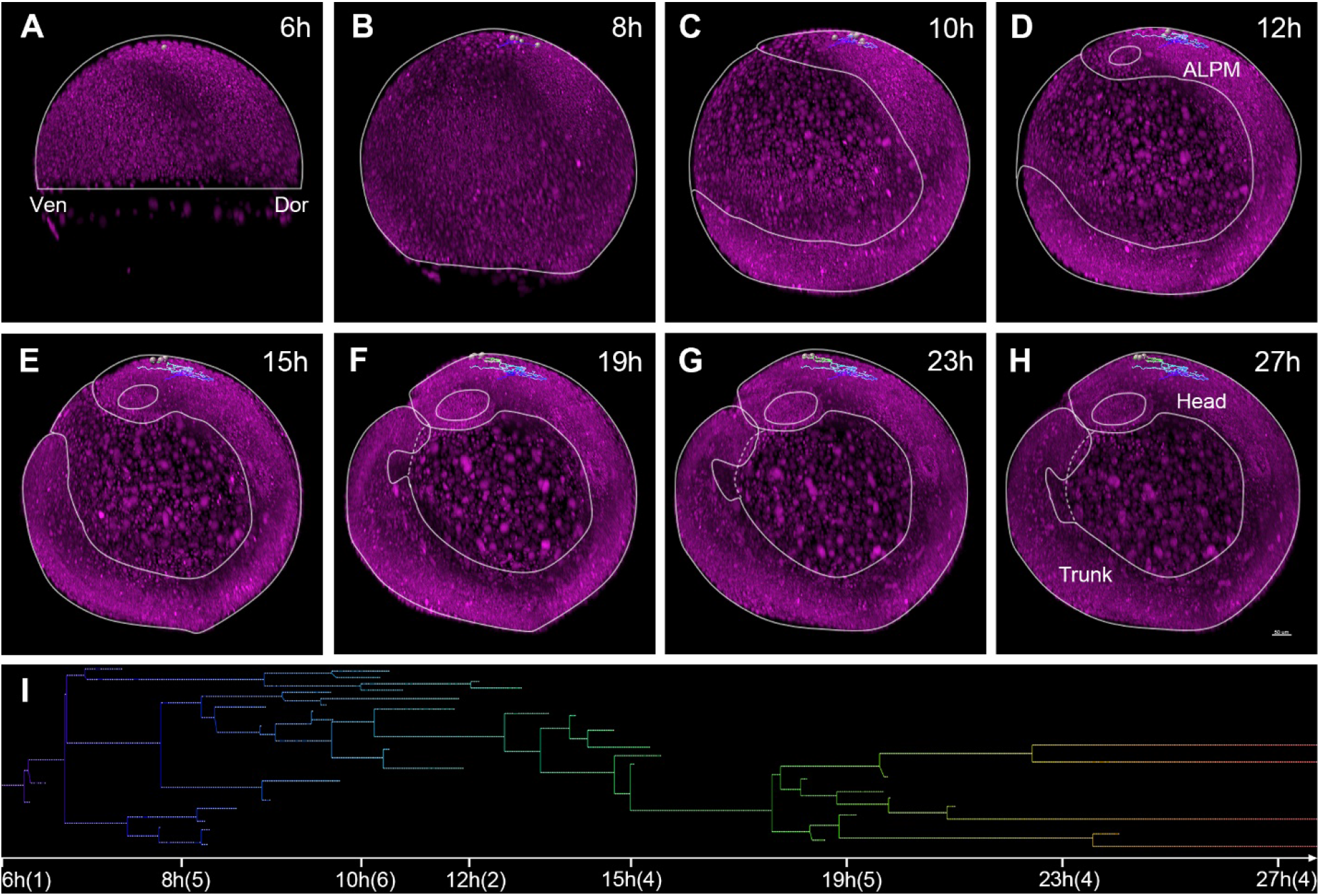
Cell-lineage tracing of early zebrafish embryos at single-cell resolution. (A, B) A selected cell near the animal pole at 6 hpf (A) divides and migrates upwards to the animal pole, and then migrates to the right side of the embryo at ~8 hpf (B). (C, D) These descendants begin to migrate back at 9 hpf, a group of descendants disappears, and the remaining group continues to migrate to the upper left of the starting position at 12 hpf. (E–H) These descendants migrate further and finally reach the top of the head between the eyes (dynamic lineage tracking in Supplementary Movie S16). (I) The cell-lineage tree map from a single gastrula cell in the animal pole at 6 hpf to four head endothelial cells at 27 hpf; note that some descendants disappear during development. Scale bar, 50 μm.

**Supplementary Fig. S8.**
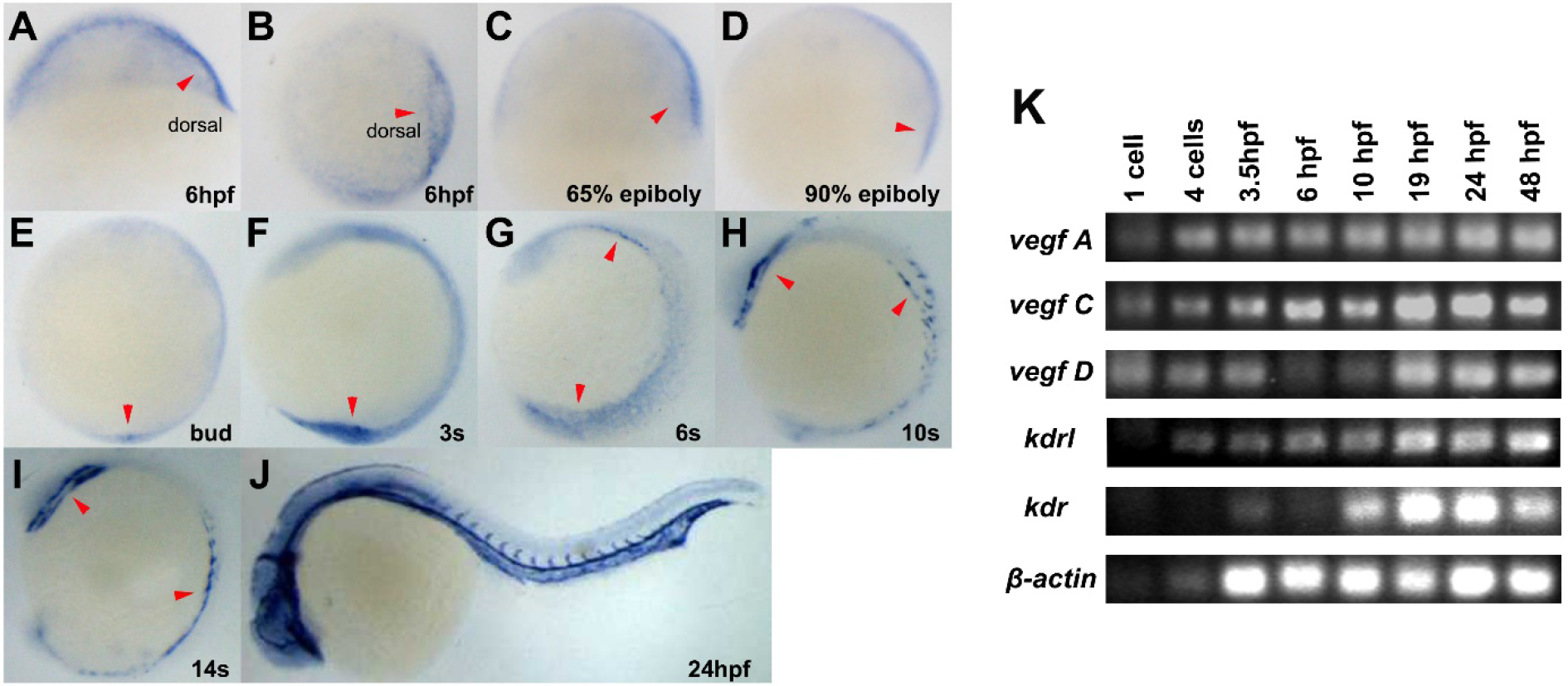
*kdrl* is enriched in the dorsal gastrula and blood vessels during early somitogenesis in zebrafish. (A–J) *In situ* RNA hybridization with the *kdrl* probe in embryos at 6 hpf (A, B), 65% epiboly (C), 90% epiboly (D), bud stage (E), 3 somites (F), 6 somites (G), 10 somites (H), 14 somites (I), and 24 hpf (J). Note the highest *kdrl* expression in the dorsal region (A, B, arrowheads), and in the ALPM and PLPM regions (G–I, arrowheads), and blood vessels (J). In A and C–J, lateral views are shown with anterior to the left. In B, top views are shown with dorsal to the right. Scale bar, 100 μm. (K) Expression of *vegfA*, *vegfC*, *vegfD*, *kdrl*, and *kdr* in 1-cell, 4-cell, 3.5-hpf, 6-hpf, 10-hpf, 19-hpf, 24-hpf, and 48-hpf embryos by semi-quantitative RT-PCR. Note that all three VEGF ligand genes (vegfA, C, and D) and the VEGF receptor kdrl had maternal expression starting from 1-cell to 48-hpf embryos, and the VEGF receptor kdr had no maternal expression starting from 3.5 to 48 hpf. β-actin was used as an internal control.

**Supplementary Fig. S9.**
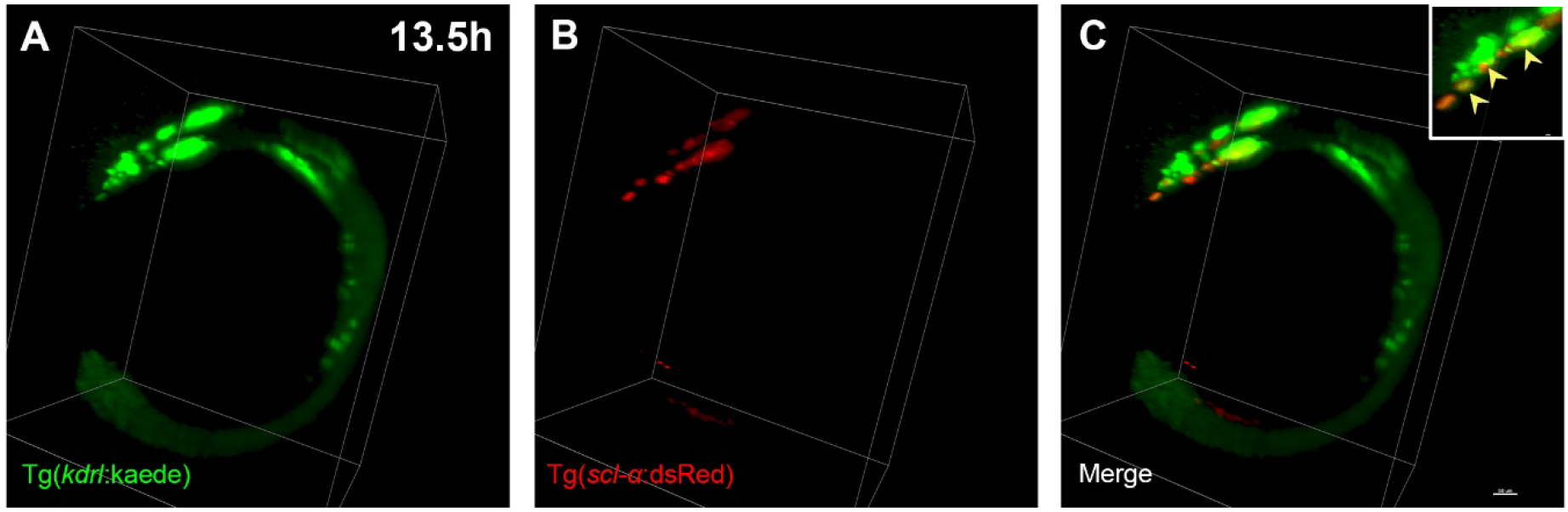
The expression of Tg(*kdrl*:kaede) is partly co-localized with Tg(*scl-α*:dsRed) in the anterior lateral plate mesoderm. (A) Image on the Tg(*kdrl*:kaede) green channel at 13.5 hpf. (B) Image on the Tg(*scl-α:*dsRed) red channel at 13.5 hpf, noting that the anterior lateral plate mesoderm was clearly labelled. (C) The merged image at 13.5 hpf. Scale bar, 50 μm.

**Supplementary Fig. S10.**
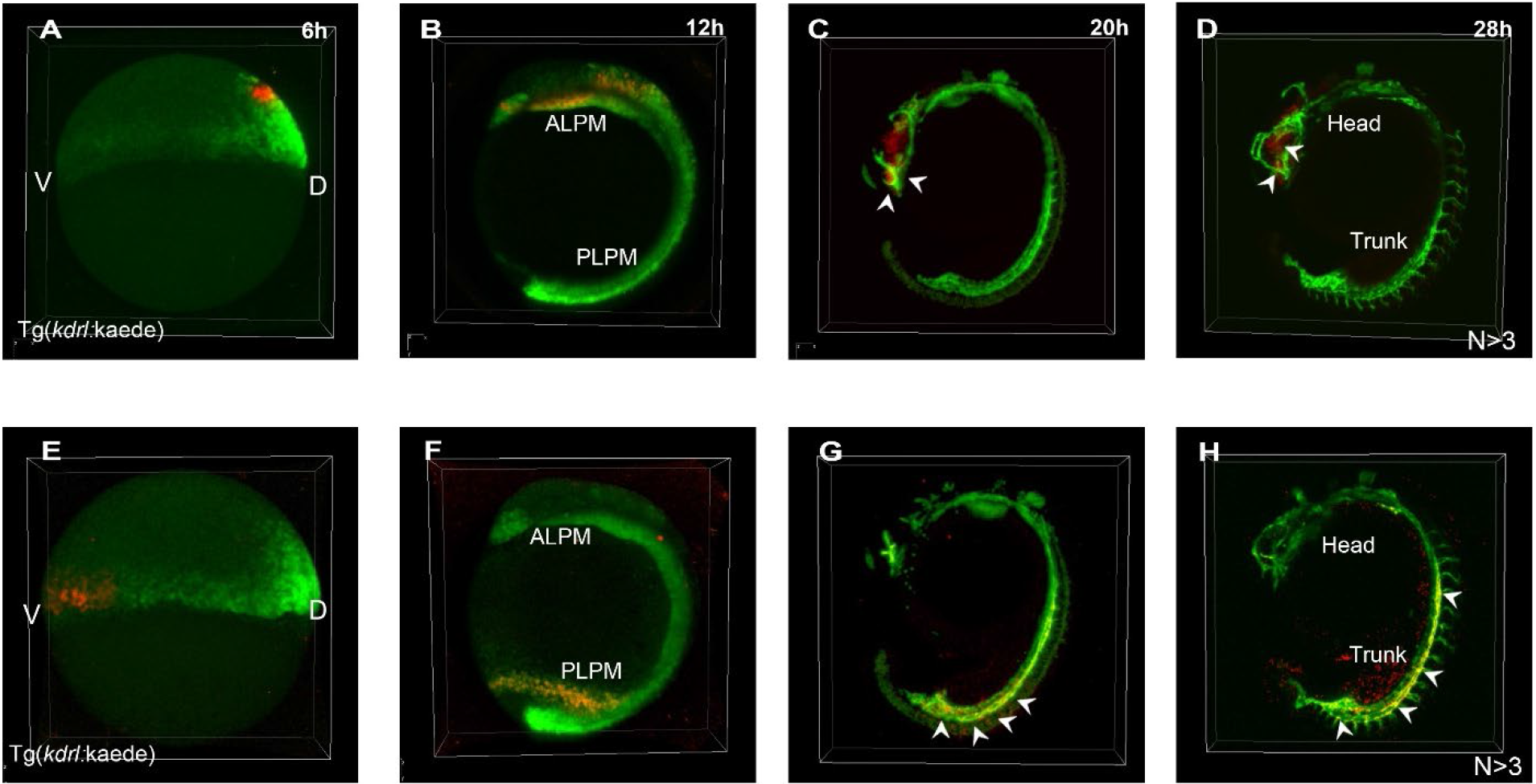
Prospective lineage tracing delineates the distinct origins of vascular ECs in the head and trunk regions. (A) Several Kaede^+^ cells in the dorsal region were converted to red by 405-nm laser excitation. (B–D) Red cells reached the anterior lateral plate mesoderm (ALPM) at 12 hpf (B) and migrated to the eyes (arrowheads) at 20–28 hpf (C, D). (E) Several Kaede^+^ cells in the ventral-lateral region were converted to red by 405-nm laser excitation. (F–H) Red cells approached the posterior lateral plate mesoderm (PLPM) at 12 hpf (F) and migrated to form trunk ECs at 20 hpf (G) and 28 hpf (H). Arrowheads point to yellow ECs; n >3.

## Supplementary Movies

The movies can be downloaded from the below website: https://disk.pku.edu.cn:443/link/F0AE2219153BAA57E90ACD6AF49E47B9

### Movie S1

Developmental morphogenesis of embryo #1 with a vegetal pole view from 6 to 27 hpf (magenta, the EGFP channel for labeling all cell nuclei); note that the embryo develops normally but is not able to stretch out, and the growing tail bends after ~19 hpf because it is embedded with FEP at an inner diameter of only 0.8 mm.

### Movie S2

Developmental morphogenesis of embryo #2 with the animal pole on the top from 6 to 22 hpf (magenta, EGFP channel for labeling all cell nuclei); note that the embryo develops normally but is not able to stretch out, and the growing tail bends after ~19 hpf, because it is embedded with FEP at an inner diameter of only 0.8 mm.

### Movie S3

Developmental morphogenesis of embryo #3 with the animal pole at the bottom from 7 to 27 hpf (magenta, EGFP channel labeling all cell nuclei); note that the embryo develops normally but isnot able to stretch out, and the tail grows under the head after ~19 hpf, because it is embedded with FEP at an inner diameter of only 0.8 mm.

### Movie S4

Representative movie showing the nuclei of embryo #1 at 6.25 hpf (magenta, nuclear EGFP signals of all cells; green, nuclear spots identified by Imaris). As the embryo rotates, the video shows an enlarged view of the nuclear EGFP signals at different locations.

### Movie S5

Representative movie showing the nuclei of embryo #1 at 16.7 hpf (magenta, nuclear EGFP signals of all cells; green, nuclear spots identified by Imaris). As the embryo rotates, the video shows an enlarged view of the nuclear EGFP signals at different locations.

### Movie S6

An embryo imaged from 6 to 20 hpf using the Luxendo Muvi-SPIM and the image dataset processed by AFEIO software (magenta, nuclear EGFP signals of all cells).

### Movie S7

Representative movie showing the maximum-intensity projections of a light-sheet time-lapse recording of embryo #1 from 6 to 27 hpf (colored lines, global cell tracks including cell movements and cell divisions; darker colors, tracks of cell positions and movements in the embryo at older time points; brighter colors, tracks and movements at younger time points). The track duration displayed is from 6 hpf to the time of recording.

### Movie S8

Retrospective lineage tracing of vascular endothelial cells in the head from 27 hpf to gastrula progenitors at 6 hpf. Representative optical images from this movie are shown in Fig. 3A–H. Magenta, nuclei; green, selected cells; colored lines, cell tracks including cell divisions and movements. The track duration displayed is from 27 hpf to the time of recording.

### Movie S9

Retrospective lineage tracing of vascular endothelial cells in the posterior trunk from 27 hpf to gastrula progenitors at 6 hpf. Representative optical images from this movie are shown in Fig. 3J–Q. Magenta, nuclei; green, selected cells; colored lines, cell tracks including cell divisions and movements. The track duration displayed is from 27 hpf to the time of recording.

### Movie S10

Retrospective fate mapping of the head vascular endothelial cells of embryo #1 from 27 hpf to dorsal-anterior gastrula progenitors at 6 hpf (red, vascular endothelial cells of the head; colored lines, retrospective tracks of vascular endothelial cells). The displayed track duration contains 100 time points.

### Movie S11

Retrospective fate mapping of anterior trunk endothelial cells of embryo #1 from 27 hpf to dorsal-lateral gastrula progenitors at 6 hpf (blue, anterior trunk endothelial cells; colored lines, retrospective tracks of vascular endothelial cells). The displayed track duration contains 100 time points.

### Movie S12

Retrospective fate mapping of posterior trunk endothelial cells of embryo #1 from 27 hpf to posterior-lateral gastrula progenitors at 6 hpf (green, posterior trunk endothelial cells; colored lines, retrospective tracks of vascular endothelial cells). The displayed track duration contains 100 time points.

### Movie S13

*In toto* imaging of the migration and division of gastrula endothelial precursors at 6 hpf to all vascular endothelial cells at 27 hpf (red, vascular endothelial cells of the head; blue, endothelial cells of the anterior trunk; green, endothelial cells of the posterior trunk; colored lines, cell movements and divisions of endothelial precursors; darker colors, cell positions and movements at older time points; brighter colors, cell positions and movements at younger time points. The track duration displayed is from 6 hpf to the time of recording.

### Movie S14

Retrospective fate mapping of vascular endothelial cells of embryo #1 from 27 hpf to gastrula progenitors at 6 hpf (red, vascular endothelial cells of the head; blue, endothelial cells of the anterior trunk; green, endothelial cells of the posterior trunk. The track duration contains 100 time points.

### Movie S15

Retrospective fate mapping of vascular endothelial cells of embryo #1 from 27 hpf to gastrula progenitors at 6 hpf (red, vascular endothelial cells of the head; blue, endothelial cells of the anterior trunk; green, endothelial cells of the posterior trunk. The track displayed is from 27 hpf to the time of recording.

### Movie S16

Developmental tracks of a single progenitor cell in the animal pole from 6 hpf to head endothelial cells at 27 hpf. Representative optical images of this movie are shown in Supplementary Fig. S7. Magenta, nuclear signals; green, selected cells; colored lines, tracks including cell movement and cell divisions; darker colors, cell positions and movements at older time points; brighter colors, cell positions and movements at younger time points. The track duration displayed is from 6 hpf to the time of recording.

